# EV-Net: A computational framework to model extracellular vesicles-mediated communication

**DOI:** 10.64898/2026.04.02.716053

**Authors:** Estefania Torrejón, Joeri Sleegers, Rune Matthiesen, Maria Paula Macedo, Anaïs Baudot, Rita Machado de Oliveira

**Affiliations:** NOVA Medical School|Universidade Nova de Lisboa, Lisbon, Portugal; Independent researcher; Aix Marseille Univ, INSERM, Marseille Medical Genetics (MMG), Marseille, France; CNRS, Marseille, France; Barcelona Supercomputing Center (BSC), Barcelona, Spain

**Keywords:** Extracellular vesicles, cell-to-cell communication, omics, networks, random walk with restart

## Abstract

**Summary:** Extracellular vesicles (EVs) are bilayer vesicles that carry a diverse cargo of molecules, such as nucleic acids, proteins and metabolites. These EVs can be transported throughout the organism to specific recipient tissues. For this reason, EVs have been recognized as pivotal mediators of cell-to-cell communication (CCC). Importantly, alterations in EV-mediated communication have been linked to pathological processes, further highlighting their biological relevance. However, the *in silico* exploration of the functional effects of EV cargo in recipient tissues remains limited due to the lack of dedicated tools that can be applied to EV omics datasets. Most current bioinformatics tools for assessing CCC rely on ligand–mediated communication and therefore cannot be used to explore EV-mediated communication.

To address this gap, we developed EV-Net, a bioinformatics tool designed to explore the effects of EV cargo on recipient tissues. EV-Net was built by adapting NicheNet, a CCC bioinformatics tool that relies on ligand-receptor mediated communication, for the analysis of EVs proteomics and RNA-seq data. The EV-Net framework enables the identification and prioritization of EV cargo molecules with high regulatory potential in a recipient tissue of interest. This prioritization facilitates the systematic translation of EV cargo profiles into testable biological hypotheses.

**Availability and documentation:** The source code of EV-Net is stored in GitHub https://github.com/torrejoNia/EV-Net alongside instructions on how to install it.

Comprehensive tutorials and additional documentation are available at https://torrejonia.github.io/EV-Net/. The datasets used in the use cases were deposited in Zenodo. The corresponding Zenodo links are provided in the tutorials for each use case. This software is distributed under a GLP3 licence.

## 1 Introduction

Multicellular life is governed by cell-to-cell communication (CCC), which enables cells to organize into tissues and organs, thereby performing more complex tasks collectively (Sgro et al., 2015). Understanding CCC is essential to uncover the molecular mechanisms underlying the pathophysiology of diseases and guide the development of new therapeutic strategies. For this reason, numerous bioinformatics tools to assess CCC have been developed, such as CellChat (Jin et al., 2021), CellPhoneDB (Efremova et al., 2020) and NicheNet (Browaeys et al., 2020). However, most of these tools rely on ligand-receptor mediated communication (Armingol et al., 2024). This approach hinders the incorporation of extracellular vesicles (EVs)-mediated communication into CCC models, as EVs uptake mechanisms in recipient tissues extend beyond classical cell surface ligand–receptor mediated communication (Mulcahy et al., 2014).

EVs are bilayer vesicles with diameters ranging from 40 to 5,000 nm that act as carriers of diverse bioactive molecules. Their EV cargo includes nucleic acids, proteins, and metabolites (Colombo et al., 2014; Kalluri et al., 2020; Kalra et al. 2016; van Niel et al., 2018). EVs are evolutionarily conserved, having been found in nearly all organisms, from unicellular to multicellular, including humans (Naito et al., 2017 and Yáñez-Mó et al., 2015). They have emerged as key mediators in CCC, contributing to the maintenance of homeostasis (Akbar et al., 2019; Huang-Doran et al., 2017; Kalra et al. 2016; Li et al., 2020). EVs mediate communication by delivering their molecular cargo to recipient cells, influencing gene expression and cellular behavior in a context-dependent manner (Van Niel et al, 2018 and Yáñez-Mó et al., 2015).

EVs are involved in a wide range of physiological and pathological processes (Naito et al., 2017; Akbar et al., 2019; Huang-Doran et al., 2017; Li et al., 2020). Dedicated bioinformatic tools to infer EV–mediated communication would be valuable, given the relevance of EVs in both healthy and pathological conditions. To the best of our knowledge, only one bioinformatic tool is currently available to assess EV-mediated CCC, miRTalk, which focuses specifically on communication driven by EVs-derived miRNAs (Shao et al., 2025).

To address this gap, we developed EV-Net, a computational framework that enables the exploration of the effect of EV cargo in their recipient tissues, using EVs proteomics or RNA-seq datasets. EV-Net was built by adapting NicheNet, an established bioinformatics tool for CCC analysis that relies on ligand-receptor mediated communication (Browaeys et al., 2020). This adaptation involved extending the NicheNet scoring algorithm so that it could be applied not only to ligands but also to other proteins that may be transported as EV cargo. We applied EV-Net to two public EVs datasets, demonstrating its ability to uncover novel insights into EV-mediated communication.

## 2 Methods

### 2.1 Adaptation of NicheNet for EV-mediated communication

EV-Net builds on NicheNet, a CCC tool that predicts ligand–target regulatory potential by integrating gene expression data with an extensive prior knowledge network of signaling and gene regulatory interactions curated from more than 15 different databases (Browaeys et al., 2020).

Originally, NicheNet was designed to analyze CCC by identifying ligands that drive gene expression changes through ligand-receptor interactions. However, EVs can modulate recipient cell function through additional mechanisms beyond ligand-receptor binding, including membrane fusion and endocytosis (Raposo and Stoorvogel 2013 and Mulcahy et al. 2014). To account for these EVs-specific modes of communication, we expanded NicheNet beyond classical cell surface ligand–receptor mediated communication. This modification enables EV cargo proteins to be linked to receptors as well as internal signaling proteins within the recipient tissue.

NicheNet utilizes a Personalized PageRank algorithm to generate a ligand-to-target scoring matrix. This algorithm involves random walk restarts using each ligand as a seed node to obtain the scoring for each ligand-target diffusion. In EV-Net, we modified this approach and applied the algorithm using not only ligands as seed nodes, but also transcription factors and intermediary signaling proteins.

### 2.2 Hyperparameters used to compute the EV cargo-to-target matrix

NicheNet was optimized by benchmarking its predictions against gene expression datasets derived from ligand perturbation experiments. This optimization process involved identifying hyperparameter values for the random walk used to compute the ligand–to–target scoring matrix, selecting those that best reproduced the transcriptional changes observed in the perturbation experiments.

As the field of EVs research is still emerging, the number of available EV cargo perturbation studies remains limited. Therefore, in EV-Net, we implemented the Personalized PageRank random walk with restart algorithm to compute the scoring matrix using the standard Nichenet hyperparameter values prior to optimization, specifically a *damping factor of 0*.*5* and an *ltf_cutoff of 0*.*99*. The *damping factor* controls the probability of restart during the random walk, thereby limiting how far signal propagation diffuses from the seed nodes. A *damping factor of 0*.*5* constrains the diffusion process, preventing the random walk from traversing overly long paths in the network. The *ltf_cutoff* determines the stringency of EV cargo–target link retention, and a value of *0*.*99* restricts the matrix to associations with the strongest regulatory potential. Overall, these modifications generated a new EV cargo-to-target matrix (Fig. 1).

**Figure 1.**
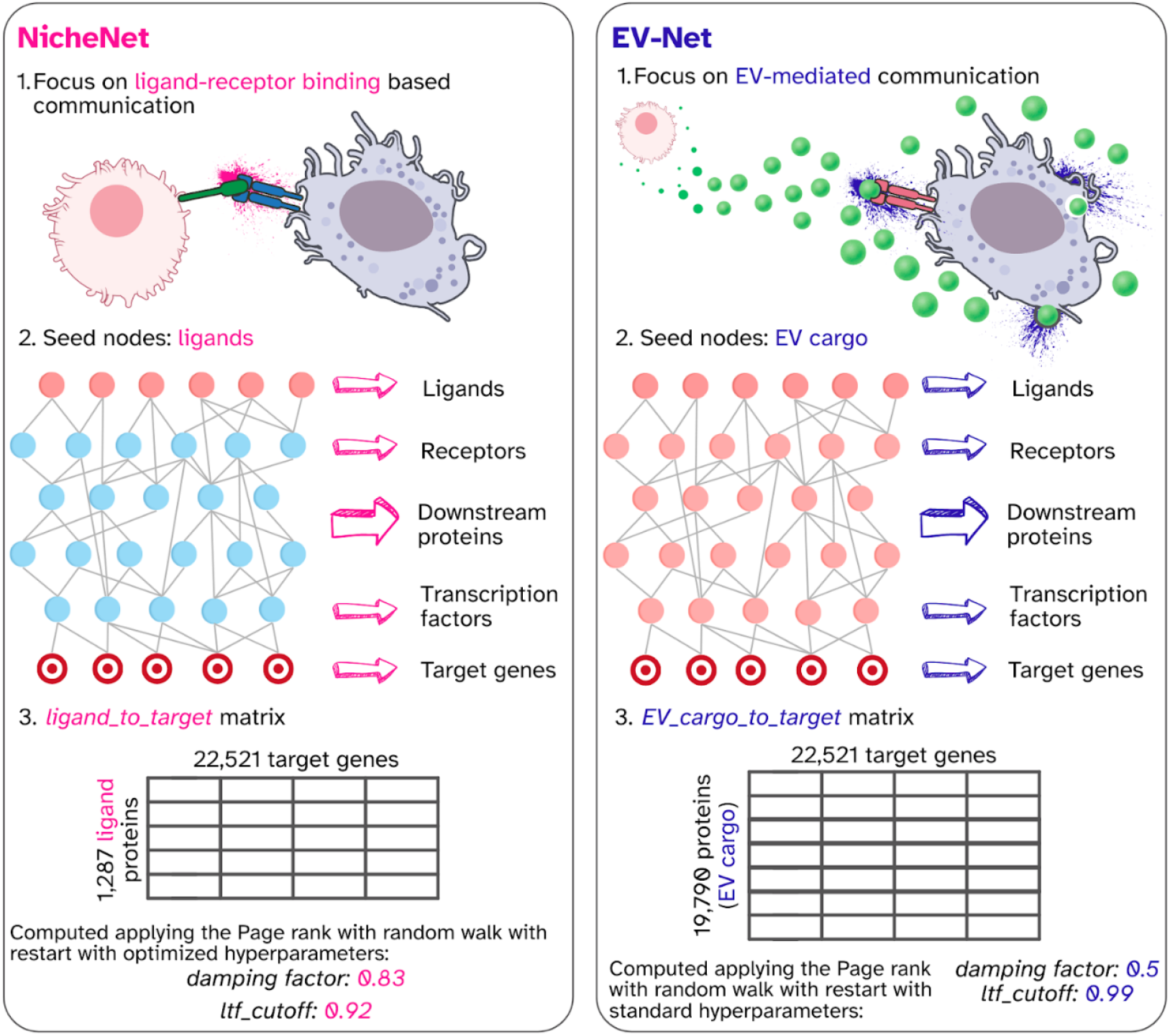
Adaptation of NicheNet for EV-mediated communication. 1.While the original NicheNet framework is specialized for ligand-receptor-based communication, EV-Net extends this paradigm to model communication mediated by the diverse molecular cargo within EVs, referred as EV cargo. 2.The Personalized PageRank algorithm is applied to the NicheNet prior-knowledge network, using ligands as seed nodes (in pink) in the case of NicheNet and EV cargo proteins in the case of EV-Net (in pink). 3.The resulting *EV cargo-to-target* matrix comprises 19,790 proteins and 22,521 target genes, replacing the original NicheNet ligand-to-target matrix, which contained 1,287 ligand proteins and 22,521 target genes.

### 2.3 Data formats

As input, EV-Net takes EV cargo data provided as a list of protein names. These may correspond to differentially abundant proteins derived from EVs proteomics datasets or to differentially expressed transcripts obtained from EVs RNA-seq datasets used as proxy for proteins. For the recipient tissue of interest, users can provide a list of differentially expressed genes derived from pre-processed single-cell or bulk transcriptomic data or a Seurat object containing expression data for the recipient tissue across two conditions or two different cell populations. When a Seurat object is supplied, differential expression analysis can be performed directly within the EV-Net pipeline. Both human and mouse datasets are supported.

As output, EV-Net generates a ranked list of EV cargo molecules with the highest regulatory potential on the target genes, defined as the EV cargo molecules that best explain the observed changes in gene expression in the recipient tissue. In addition, the framework identifies candidate target genes that are likely to be regulated by the EV cargo, offering insights into putative EV-mediated regulatory mechanisms. These results are visualized as a regulatory potential matrix, where the prioritized EV cargo are displayed along the y-axis, target genes from the recipient tissue along the x-axis, and regulatory potential is represented by color intensity (Supplementary Figures 1 and 2). Users can further select highly ranked EV cargo molecules and their associated target genes to model signaling pathways (Supplementary Figure 3).

### 2.4 Main functionalities

EV-Net assesses the effects of EV molecular cargo, including proteins or RNA transcripts, on gene expression in recipient tissues. It can compare two experimental conditions or a single condition across different cell populations. EV-Net prioritizes EV cargo molecules (derived from proteomics or RNA-seq transcripts used as protein proxies) by ranking them according to their predicted regulatory potential on target gene expression. This process allows for the inference of putative EV cargo-target gene links specific to the recipient tissue under study.

When comparing two conditions, the user needs to provide the differentially abundant proteins or differentially expressed RNA-seq transcripts present in EVs as the EV cargo for each condition, together with the corresponding differentially expressed genes in the recipient tissue. The tutorials provided on our website and in the repository offer guidance for performing these analyses.

## 3 Use-cases

We demonstrate the utility of EV-Net by exploring the effect of EVs in CCC using EV cargo proteomic data in two different scenarios.

### 3.1 Effect of gut EVs in Kupffer cells

We applied EV-Net to investigate the impact of gut-derived EV cargo from a prediabetic mouse model on the liver-resident macrophages, Kupffer cells, as the recipient tissue of interest.

To inform our selection of EV cargo, we used proteomics data from gut-derived EVs isolated from prediabetic and healthy mice (Ferreira et al., 2022). We selected EVs proteins after a differential abundance analysis by an empirical Bayes moderated t-test implemented in limma (Ritchie et al., 2015). Specifically, proteins with a *p-value < 0*.*1* and a *log fold change > 1* corresponding to the most abundant in the prediabetic condition.

We chose Kupffer cells as the recipient tissue of interest due to their role in liver metabolism, inflammation and lipid remodeling (Barreby et al., 2022). We focused on genes that are upregulated in Kupffer cells compared to hepatocytes using single-cell transcriptomic data from liver cells obtained from the Tabula Muris Atlas (Tabula Muris Consortium, 2018). The differential expression analysis between these cell populations was performed using the Seurat function FindMarkers (Butler et al., 2028) and filtered by an adjusted *p-value <= 0*.*05* and *log fold change >= 0*.*25*. This approach enabled us to prioritize the EV cargo with high regulatory potential on Kupffer cells–enriched genes.

Using this approach, we identified the selenoprotein Selenocysteine Lyase (SCLY) among the EV cargo proteins showing high regulatory potential on numerous Kupffer cell genes (Supplementary Fig. 1). SCLY is involved in oxidative stress management by providing an alternative source of selenium (Labunskyy et al., 2014). Selenium plays a role in maintaining redox balance and attenuating the pro-inflammatory activity of Kupffer cells (Shilo et al., 2008).

This finding suggests that gut-derived EVs-associated SCLY may contribute to the modulation of Kupffer cell function in a prediabetic context.

### 3.2 Effect of lipopolysaccharide treated microglia-derived EVs on recipient naïve microglia

As a second use case, we investigated the effect of EVs released by lipopolysaccharide (LPS)-activated microglia on unstimulated (naïve) microglia. LPS induces a pro-inflammatory phenotype in microglia, the brain’s resident immune cells, which play a central role in mediating neuroinflammatory responses across various neurological disorders (Santiago et al., 2023).

We used publicly available proteomics data from EVs derived from LPS-activated microglia to study the EV cargo (Santiago et al., 2023). In this study, Santiago and colleagues performed a differential abundance analysis using unpaired *t*-tests to compare EVs proteomics data from control microglia vs. LPS-activated microglia and other experimental conditions. Proteins that were more abundant in EVs from LPS-activated microglia and showed *p-values < 0*.*1* were selected as EV cargo for our analysis.

As input data for the recipient tissue, we utilized transcriptomics data from naïve microglia exposed to LPS-activated microglia-derived EVs provided in the same study by Santiago et al., 2023. We focused on naïve microglia genes that were differentially expressed in the condition treated with LPS-activated microglia-derived EVs, based on an *adjusted p-value < 0*.*1* and an *absolute log fold change >= 0*.*25*.

Santiago and colleagues reported that naïve microglia adopted a pro-inflammatory phenotype following exposure to the LPS-activated microglia-derived EVs. Using EV-Net, we identified the protein metadherin (MTDH) as an EV cargo molecule with high regulatory potential on inflammation-related genes in microglia, such as discoidin domain receptor 1, DDR1 (Supplementary Fig. 2). Although MTDH is known as an oncogenic protein in several cancers, its role in microglia biology has not been previously explored. These findings indicate that EVs-associated MTDH may act as a mediator contributing to inflammatory activation of microglia.

## 4 Discussion

To date, more than 4,000 studies have contributed EV-related datasets to public repositories like ExoCarta (Keerthikumar et al., 2015), Vesiclepedia (Chitti et al., 2023) and EV-COMM (Chen et al., 2024). The growing availability of EV cargo datasets represents an opportunity to further explore the role of EVs in CCC. However, determining the biological impact of EVs on recipient tissues remains a major challenge in biomedical research. While EV cargo profiling has become increasingly comprehensive, translating complex proteomic or transcriptomic signatures into mechanistic insight and experimental hypotheses is still largely non-trivial. This gap between data generation and biological interpretation often hampers the design of functional studies and slows progress toward understanding EV-mediated communication. In this context, EV-Net was developed as a computational framework to bridge EVs omics data with recipient tissue. Our framework enables the prioritization of biologically relevant EV cargo and the exploration of the effects of said cargo in the recipient tissue. Overall, EV-Net supports the generation of hypotheses by the identification of EV cargo-target genes links that were previously unknown or not taken into consideration in the recipient tissue under study. EV-Net facilitates targeted and more efficient downstream *in vitro* and *in vivo* experimentation.

EV-Net represents one of the first computational frameworks specifically designed to explore the effect of EV cargo in their recipient tissues. Other approaches are beginning to emerge, such as miRTalk, which focuses specifically on EVs-derived miRNA-mediated communication (Shao et al., 2025). To our knowledge, EV-Net is the first framework to leverage EVs proteomics and RNA-seq data for the exploration of EV-mediated communication.

EV-Net builds on the NicheNet framework, granting access to an extensive prior knowledge network curated from more than 15 databases encompassing protein–protein and gene regulatory interactions (Browaeys et al., 2020). While this is a major strength, it also introduces certain limitations. In particular, interactions in the NicheNet prior knowledge network are not annotated as activating or inhibitory (Browaeys et al., 2020). Consequently, EV-Net primarily predicts activating signaling links, as the underlying network lacks the directional annotations required to explicitly infer inhibitory signaling.

The EVs field is still rapidly evolving, and several important biological questions remain unresolved. For instance, the stoichiometric fraction of EV cargo that remains functional upon reaching the recipient tissue cytosol is not yet well-defined. (O’Brien et al., 2022 and Ripoll et al., 2026). The current version of EV-Net operates under the assumption that all identified cargo molecules possess the potential to regulate the recipient tissue. In reality, the proportion of active EV cargo may vary. This proportion could potentially be estimated based on the abundance of specific ligands within EVs, which would influence processes such as receptor binding, endocytosis, or membrane fusion (Mulcahy et al. 2014; Ginini et al., 2022 and Bonsergent et al., 2021). These aspects were not incorporated in the design of EV-Net, highlighting areas where the framework could be further refined as more experimental data becomes available.

Ultimately, EV-Net translates EVs molecular profiles into mechanistic hypotheses, ranks EV cargo molecules, and guides functional experimental design. By providing a bridge between EVs omics data and biological interpretation, EV-Net offers a powerful strategy to advance both fundamental EVs biology and its translational applications in biomedicine.

## Author contributions statement

Conceptualization: RMO, AB and ET

Funding acquisition: RMO, AB, MPM and ET

Methodology: AB and ET

Software: ET, JS and RM

Supervision: RMO, AB, MPM and RM

Writing: RMO, AB, ET, JS and RM

Validation: ET, JS and RM

Visualization: ET

## Data availability

EV-Net can be installed from the GitHub repository https://github.com/torrejoNia/EV-Net. Documentation, together with two tutorials, are hosted at the repository website https://torrejonia.github.io/EV-Net/. This website was created using Quarto (Allaire et al., 2026).

The datasets used in the use cases were deposited in Zenodo. The corresponding Zenodo links are provided in the tutorials for each use case.

## Acknowledgments

We thank the members of the Systems Biomedicine Lab at Marseille Medical Genetics for the valuable discussions that supported this project. In particular, we thank Céline Chevalier and Benjamin Loire for their technical support, and Morgane Térézol and Bastien Chassagnol for their recommendations regarding documentation packages. We also thank Pedro Ribeiro from the MEDIR Lab at NOVA Medical School for testing the tool. Finally, we acknowledge Yvan Saeys’s team, the creators of NicheNet, for their feedback on our tool.

