## Supplementary figures for "EV-Net: A computational framework to model extracellular vesicles-mediated communication"

Figure 1. Prioritized EV cargo derived from gut and their regulatory potential in Kupffer cells target genes.

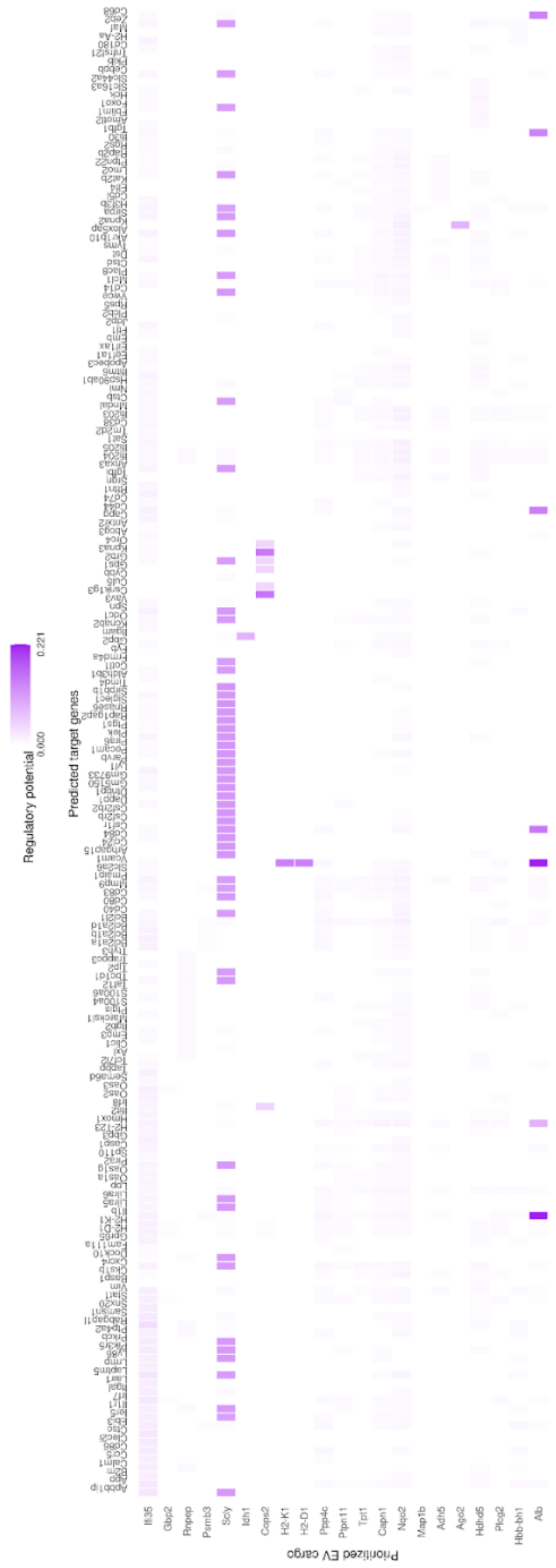

Figure 2. Prioritized EV cargo derived from LPS-activated microglia and their regulatory potential in naïve microglia target genes.

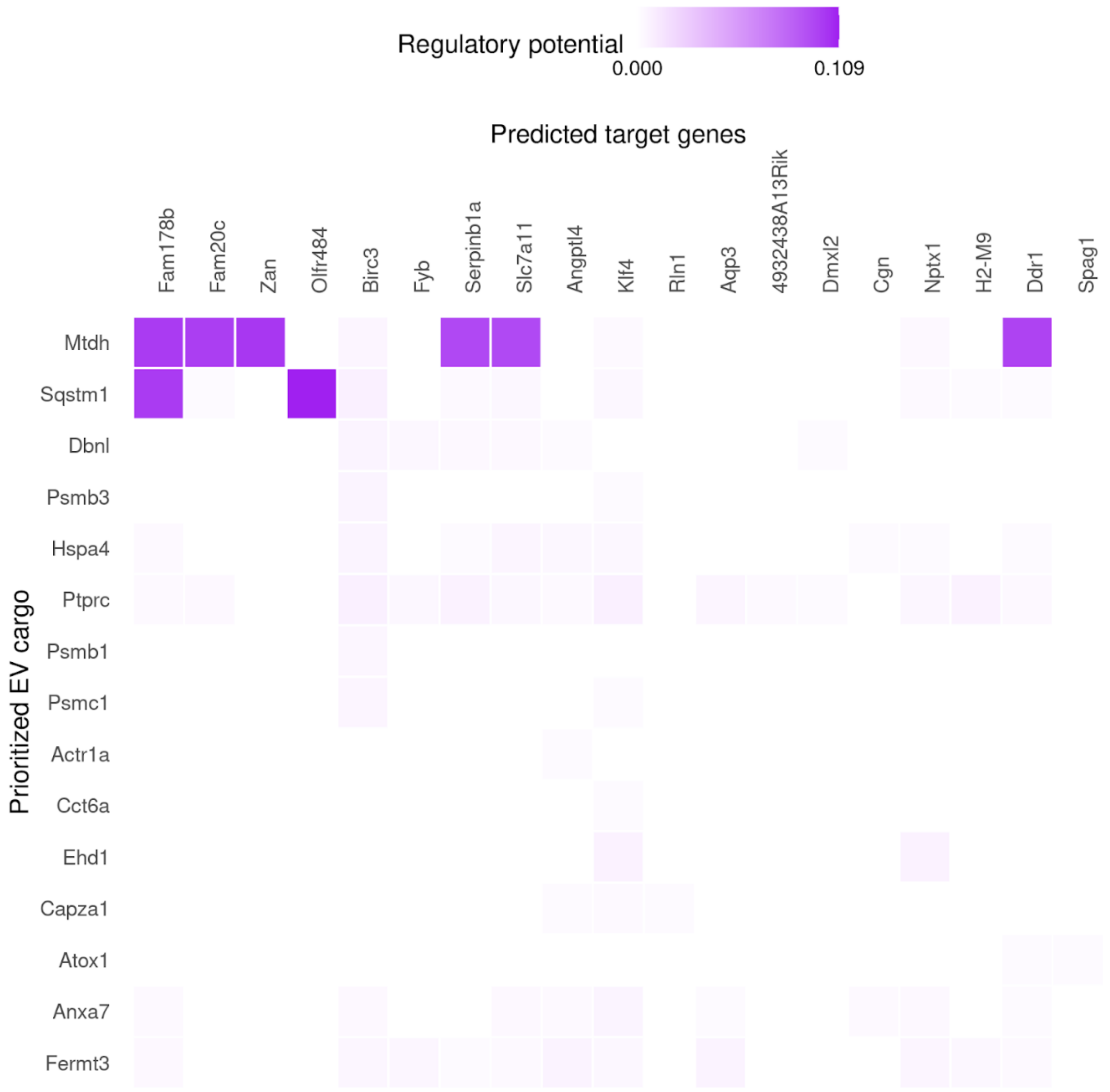


Figure 3. Signaling paths from the gut-derived EV cargo protein “SCLY” to possible Kupffer cells target genes “ZEB2” and “FOXO1”

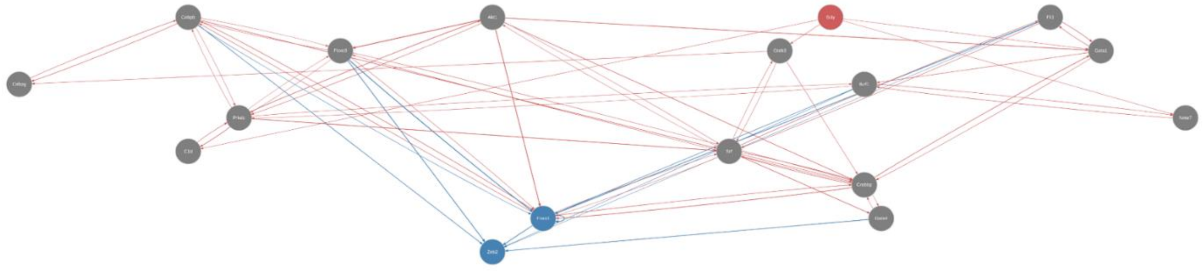
